# A metagenomic analysis of the wrackbed microbiome indicates a phylogeographic break along the North Sea - Baltic Sea transition zone

**DOI:** 10.1101/2021.11.01.466799

**Authors:** Emma L. Berdan, Fabian Roger, Alexandra Kinnby, Gunnar Cervin, Ricardo Pereyra, Mats Töpel, Maren Wellenreuther, Kerstin Johannesson, Roger K. Butlin, Carl André

## Abstract

Sandy beaches are biogeochemical hotspots that bridge marine and terrestrial ecosystems via the transfer of marine organic matter, such as seaweed (termed wrack). A keystone of this unique ecosystem is the microbial community, which helps to degrade wrack and re-mineralize nutrients. However, little is known about the wrackbed microbiome, its composition, trophic ecology, or how it varies over time and space. Here we characterize the wrackbed microbiome as well as the microbiome of a primary consumer, the seaweed fly *Coelopa frigida*, and examine how they change along one of the most studied ecological gradients in the world, the transition from the marine North Sea to the brackish Baltic Sea. We found that polysaccharide degraders dominated both the wrackbed and seaweed fly microbiomes but there were still consistent differences between wrackbed and fly samples. Furthermore, we observed a shift in both microbial communities and functionality between the North and Baltic Sea. These shifts were mostly due to changes in the frequency of different groups of known polysaccharide degraders (Proteobacteria and Bacteroidota). We hypothesize that microbes were selected for their abilities to degrade different polysaccharides corresponding to a shift in polysaccharide content in the seaweed communities of the North vs. Baltic Sea. Our results reveal the complexities of both the wracked microbial community, with different groups specialized to different roles, and the cascading trophic consequences of shifts in the near shore algal community.

## Introduction

Sandy beaches comprise 31% of the world’s ice free coastline (Luijendijk, Hagenaars et al. 2018) and represent some of the most ecologically and economically valuable landforms (Barbier, Hacker et al. 2011). Beaches bridge marine and terrestrial ecosystems and provide critical ecosystem functions to both, such as recycling nutrients (Koop, Newell et al. 1982, Rodil, Lastra et al. 2019, Hyndes, Berdan et al. 2021) and supporting key habitats, for example bird nesting sites (Schlacher, Hutton et al. 2017).

Unlike other crucial ecosystems, beaches have little to no primary productivity (Colombini and Chelazzi 2003). Instead, organic matter, such as algae and carrion, is deposited on the beaches forming the basis of the sandy beach ecosystem (Colombini and Chelazzi 2003, Hyndes, Berdan et al. 2021). These deposits, called wrackbeds, are primarily decomposed by bacteria, physical processing, and invertebrate consumption (Jêdrzejczak 2002, Colombini and Chelazzi 2003, Lastra, Rodil et al. 2014, Rodil, Lastra et al. 2019, Hyndes, Berdan et al. 2021). Wrackbeds are biogeochemical hotspots with extremely high metabolic activity (Rodil, Lastra et al. 2019) partly due to bacteria; after algae are deposited on the beach, bacterial densities increase up to four orders of magnitude (Koop, Newell et al. 1982, Cullen, Young et al. 1987, Urban-Malinga and Burska 2009). Along with detritivores, these bacteria mineralize nutrients, which are then exported back to the sea (Koop, Newell et al. 1982, Dugan, Hubbard et al. 2011, Rodil, Lastra et al. 2019, van Erk, Meier et al. 2020). This microbial biomass then serves as the basis for secondary production, providing food for macro and meiofauna, such as dipteran larvae, nematodes, and amphipods (Cullen, Young et al. 1987, Urban-Malinga and Burska 2009, Porri, Hill et al. 2011, Griffin, Day et al. 2018, Singh, Huggett et al. 2021). Secondary consumers, in turn, such as spiders and beetles, and scavengers such as birds and mammals then prey on the macro and meiofauna (Hyndes, Berdan et al. 2021). As such, wrackbed-decomposing microbiota form the basis of the beach ecosystem. However, we understand little about these communities, their composition, their functionality, and how they vary over space and time. We also know little about the ecological consequences of variation in the wrackbed microbiome (e.g., the bacterial species composition), and its effects on associated species. Changes in the wrackbed microbiome likely affect the composition of the eukaryotic consumer community by exerting different selective pressures on individual species, similar to the interaction between soil microbiomes and plant communities (Trivedi, Leach et al. 2020).

One such consumer group and common inhabitant of wrackbeds are seaweed flies, such as *Coelopa frigida* in Northern Europe. Eggs of this species are laid on the seaweed and emerging larvae feed primarily on the wrackbed microbiome (Cullen, Young et al. 1987). The flies experience high mortality during the early larval phase (Butlin and Day 1984, Cullen, Young et al. 1987), with mortality and growth rates differing based on the seaweed composition within the wrackbed (Cullen, Young et al. 1987, Edward 2008). Data indicate that shifts in the seaweed composition of the wrackbed are accompanied by shifts in the genetic composition of inhabiting seaweed fly populations, notably in the frequency of the *Cf-Inv(1)* chromosomal inversion (Day, Dawe et al. 1983, Butlin and Day 1989, Wellenreuther, Rosenquist et al. 2017, Berdan, Rosenquist et al. 2018, Mérot, Berdan et al. 2018). This suggests that the wrackbed microbiome could exert significant selective pressure on both *C. frigida* and even on the *Cf-Inv(1)* inversion itself (Edward 2008, Edward and Gilburn 2013), pointing towards a potential importance of the wrackbed microbiome for higher trophic levels. However, whether or not the wrackbed microbiome is one driver of selection in this species remains unknown.

Here, we examine the structure and function of the wrackbed microbiome along one of the most studied ecological gradients in the world, the transition from the marine North Sea to the brackish Baltic Sea. Multiple abiotic factors vary along this transition zone, including salinity, temperature, and alkalinity (Møller Nielsen, Paulino et al. 2016, Snoeijs-Leijonmalm, Schubert et al. 2017). This is accompanied by shifts in algal, seagrass, and seawater microbial communities (Herlemann, Labrenz et al. 2011, Schubert, Feuerpfeil et al. 2011, Herlemann, Lundin et al. 2016, Takolander, Cabeza et al. 2017). We sampled wrackbeds from five sites spanning the transition and investigated how the species and functional composition of the bacterial communities changed over this gradient. In a second step, to understand how changes in the wrackbed microbiome composition impact the food chain, we sequenced the microbiome of seaweed fly larvae from the same sites. We used these data to ask three key questions: 1. How does the species composition and functionality of the wrackbed microbiome vary over the transition zone?, 2. Can these changes be linked to specific environmental factors?, and 3. Can we detect cascading effects of changing microbiome community composition on seaweed flies?

## Methods

### Sample Collection

Samples were collected in July and August 2016 from five sites along the Scandinavian Coastline: Skeie (58°41’50.4”N 5°32’27.0”E) and Justoya (58°13’08.2”N 8°23’12.1”E) in Norway and Magnarp (56°17’51.0”N 12°47’18.4”E), Smygehuk (55°20’17.3”N 13°21’48.7”E), and Ystad (55°25’27.9”N 13°46’23.1”E) in Sweden. We split these sites into two Baltic and three non-Baltic sites (Figure 1; Snoeijs-Leijonmalm and Andrén 2017).

**Figure 1.**
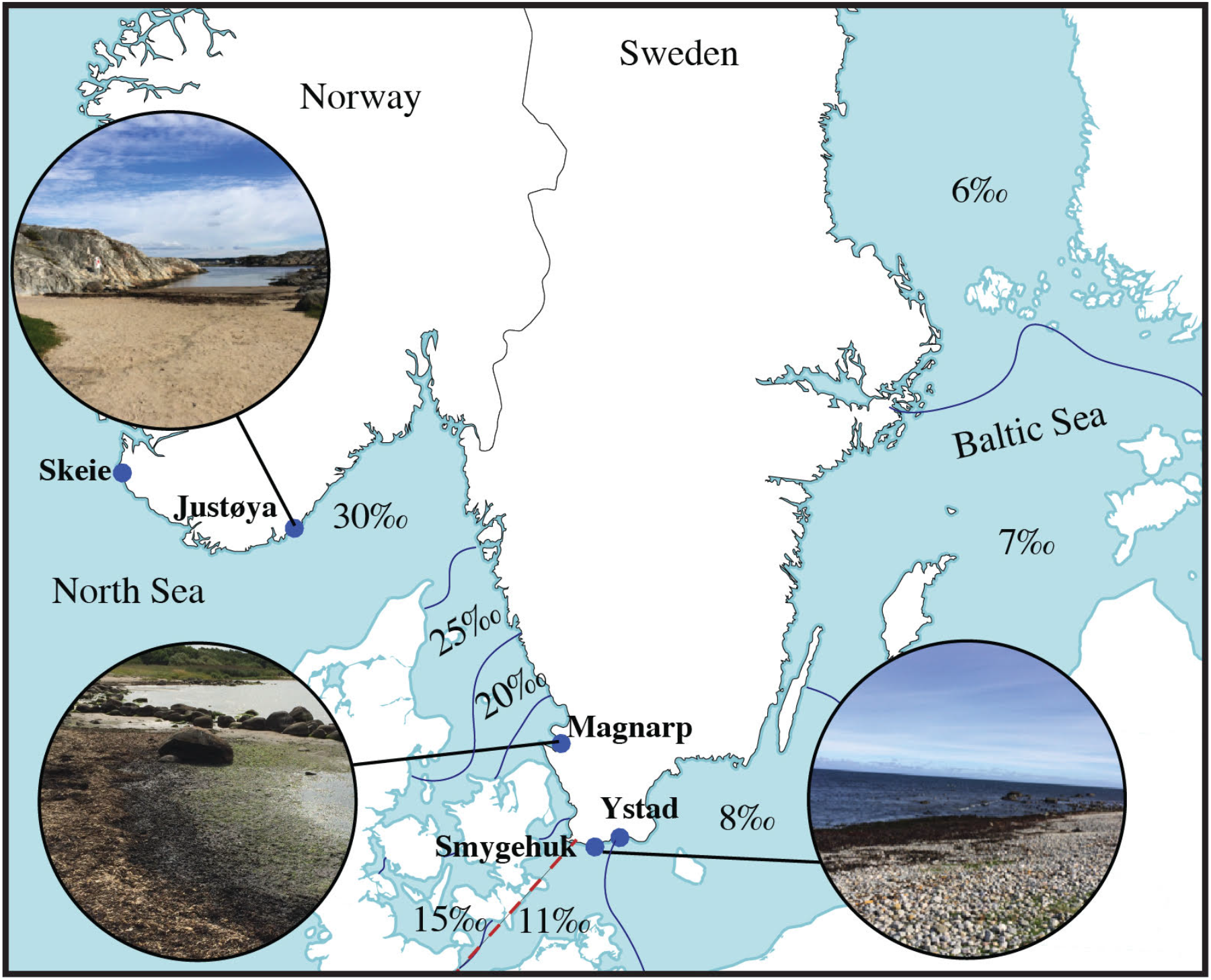
Map of sampling sites with representative photos of wrackbeds from some sites as well as the salinity gradient from the North Sea into the Baltic Sea. The red dashed line indicates the split between Baltic and non-Baltic sites.

We collected both wrackbed and larval samples from each site. We collected handfuls of seaweed from widely spaced parts of the wrackbed where *C. frigida* larval density was high (more than 50 larvae in approx. a handful of wrackbed). For three of these handfuls we removed as many larvae as possible and then placed the remaining matter in a 50 ml tube filled with 99% ethanol. All other handfuls were placed in 2-3 ventilated plastic containers. After collection, we chose 20 random larvae per wrackbed which were placed in groups of 5 in 1.5 ml Eppendorf tubes filled with 99% ethanol. Fresh seaweed samples were also collected from the tideline to gage the seaweed compositional make-up of the site (with the exception of Justøya). All samples were transported back to Tjärnö in Sweden, where they were stored at −20°C until processing.

### DNA Extraction and Library Preparation

We separately extracted DNA from wrackbed samples and individual larvae. All remaining larvae were removed from the wrackbed samples, the wrackbed material was spun down for 10 minutes at 3220 rcf, and excess ethanol was poured off. The samples were flash frozen using liquid nitrogen and subsequently ground with a mortar and pestle. Two technical replicates of each wrackbed sample (6 extractions per wrackbed) were extracted using the MoBio PowerSoil DNA Isolation Kit (Carlsbad, CA) following the manufacturer’s instructions. Individual larvae were removed and allowed to dry before being extracted in the same manner. We extracted 15-17 larvae per site.

To examine the relationship between the microbiome and the genetic structure in *C. frigida* we genotyped all larvae for the *Cf-Inv(1)* inversion. In *C. frigida*, frequencies of the *Cf-Inv(1)* inversion vary, depending on a variety of abiotic and biotic factors, including the seaweed composition of the wrackbed (Day, Dawe et al. 1983, Butlin and Day 1989, Mérot, Berdan et al. 2018). Thus, we hypothesized that there may be a relationship between larval microbiome and *Cf-Inv(1)* genotype. Larvae were genotyped for the *Cf-Inv(1)* inversion using a diagnostic SNP as described in Mérot et al. (2018).

To examine the wrackbed microbiome we used amplicon sequencing targeting the V3–V4 loops of the bacterial and archaeal 16S genes with the 341F and 805R primers. We followed the protocol from the Andersson lab (https://github.com/EnvGen/LabProtocols/blob/master/Amplicon_dual_index_prep_EnvGen.rst) to generate individually-barcoded libraries for each of our samples (4 wrackbed samples (including 1 technical replicate) and 15-17 larvae per site). Samples were then pooled and sequenced on one flowcell of MiSeq v3 (paired-end 300 bp reads) at the National Genomics Infrastructure in Stockholm, Sweden.

### Data Processing

After de-multiplexing, primer and adaptor removal, as well as trimming, were done using cutadapt (Martin 2011). We assembled these quality-filtered reads into error-corrected Amplicon Sequence Variants (ASVs) (Callahan, McMurdie et al. 2017), using DADA2 v1.18.0 (Callahan, McMurdie et al. 2016), largely following the DADA2 pipeline tutorial. In brief, read quality of primer-trimmed forward and reverse reads was visualized and after manual inspections of the profiles we chose a truncation parameter of 270 bp for the forward reads and 200 bp for the reverse reads – ensuring an overlap of 45 base-pairs. During quality filtering we allowed for 2 (forward) and 5 (reverse) expected errors after trimming. This setting resulted in a median loss of 40% of the reads (inter-quartile range 37 - 45.5%). De-replication, de-noising and merging of the paired reads were performed using default parameters, choosing the “pseudo” option for de-noising. After merging, sequences with a length of greater than 431 bp or shorter than 399 bp were discarded. This excluded 30% of the ASVs accounting for 2.7% of the (remaining) reads. We checked for chimeric sequences using the “consensus” method from the removeBimeraDenovo function and all sequences identified as likely chimeras were discarded (30% of ASVs, 1% of the reads). Assembled ASVs were assigned a taxonomy using the Ribosomal Database Project (RDP) naïve classifier method (Wang, Garrity et al. 2007) implemented in the assignTaxonomy function in DADA2 - using the The SILVA ribosomal RNA gene database v138.1 (Quast, Pruesse et al. 2012) as reference. Taxonomic assignment at any rank was only maintained if the taxon was assigned a probability of >= 80% (default setting) by the RDP classifier. Reads that were not classified at Kingdom level or were classified as one of Eukaryota, Chloropast or as Mitochondria were discarded (2% of ASVs, 0.7% of the reads). As an additional quality-filtering step, we aligned all sequences using the AlignSeqs function from the DECIPHER R package (Wright 2016) and calculated the sum of the distance of each sequence to all other sequences. Visual inspection of the distribution of distance-sums revealed a group of sequences that was almost twice as different from all other sequences and the vast majority of these sequences had no taxonomic annotation at phylum level. This makes it likely that these ASVs represent non-biological sequences (e.g., undetected chimeras) and therefore we excluded them (2% of ASVs, 0.07% of the reads).

The stringent quality filtering described above resulted in an ASV table with 13,125 unique ASVs across all samples. As a final step we clustered the sequences to operational taxonomical units (OTUs) at 99% identity using vsearch v2.17.0 (Rognes, Flouri et al. 2016) to reduce any remaining spurious diversity and because our goal was to compare the community composition across sites and not to study any specific strains. The clustering resulted in 7,775 unique OTUs. OTUs present in only a single sample or with less than 5 reads across all samples were further excluded (40% of the ASVs, 0.8% of the sequences), resulting in a final OTU table with 4,655 OTUs.

Our data set contained five pairs of technical replicates from the wrackbeds. To test our pipeline, we visualized all wrackbed samples in an NMDS plot with Bray-Curtis dissimilarity using the package ‘phyloseq’ (McMurdie and Holmes 2013). All technical replicates were closer to their partner than to any other sample with the exception of a single sample from Skeie that appeared to have been mislabeled (Figure S1). We removed one technical replicate from each pair (the one with fewer summed ASV counts) and the anomalous Skeie sample.

### Statistical Analysis

We built a phylogenetic tree of all OTU sequences for use in downstream analyses. We used the DECIPHER R package (Wright 2016) to create a multiple-alignment of all of our sequences. We then staggered our alignments using the StaggerAlignment function and built an approximate-maximum-likelihood tree using FastTree (v2.1.11), then used the phangorn R package (Schliep 2010) to construct a neighbor joining tree. Using this as a starting point we then fitted a maximum likelihood tree assuming the GTR+G+I mutation model (Generalized Time-Reversible with Gamma rate variation).

This tree was used with the PhILR R package (Silverman, Washburne et al. 2017) to perform a Phylogenetic Isometric Log-Ratio Transformation on our data (Silverman, Washburne et al. 2017). This is a compositionally aware approach that controls for false positives by testing for the changes in log ratios between microbial abundances (called balances) that are constructed using evolutionary history (i.e., the phylogenetic tree). This technique fully accounts for the correlation structure of the data as well as the compositional nature of the data (Gloor, Macklaim et al. 2017). We performed ordinations with Euclidian distance using the phyloseq package (McMurdie and Holmes 2013) which indicated a difference between Baltic and non-Baltic samples. We identified balances that separated Baltic and non-Baltic samples using a sparse logistic regression from the glmnet package (Friedman, Hastie et al. 2010) implemented in R. The lambda penalty for this regression was estimated using our data and the cv.glmnet function. We extracted the PhiLR Euclidian distance using the vegdist function implemented in the vegan package (Dixon 2003). We then performed a multivariate ANOVA to determine which factors separated the samples using the adonis function (Dixon 2003).

To examine the functional structure of our data we estimated the potential functional roles of OTUs using the Functional Annotation of Prokaryotic Taxa (FAPROTAX) database v1.2.4 and following the method of Louca et al. (2016). We were able to assign at least one function to 1,136 of our 4,655 OTUs (24%). Overall, 69 functional groupings were associated with at least one OTU but we removed all groupings associated with fewer than 3 OTUs (14 groups) and groups that had a similarity of 1 (Jaccard similarity index) with one other group (9 groups). We used these functional data to perform an ordination in phyloseq using Bray-Curtis distances. Then, using the vegan package in R (Oksanen, Kindt et al. 2008), we performed a PERMDISP analysis to determine if different groups were differently dispersed. To examine the relative abundance of the different functions, we further removed groups that had a Jaccard similarity index of >0.75 with another group.

## Results

Sampled microbiomes were highly site-specific and variable across 15 wrackbed and 74 larval samples from five sites along the North Sea to the Baltic Sea transition zone. Seaweed composition at these sites was also highly variable (Table S1). We identified 4,655 OTUs, most of which were present in only a subset of samples (Figure S2); mean prevalence was 0.141 ± 0.002 across the 89 samples, and there were no OTUs present in all samples. Each sample contained between 52 and 1,691 OTUs (median - 610, interquartile range - 651; full sample information can be found in Table S2). The distribution of OTUs across sites was non-random, only 13.9% of OTUs (646) were found in at least one sample at all five sites, whereas 28% (1,304) were only found at a single site. Some OTUs were sample type specific; 10.1% of OTUs (472) were unique to wrackbed samples and 1.4% (66) were unique to larvae. Wrack samples grouped strongly by site (Figure S3) but larval samples overlapped somewhat. Rarefaction curves showed that most samples were asymptotic (Figure S4) indicating that further sequencing effort would not greatly affect the results. To look at diversity in our samples we calculated the effective number of species of order q = 1. The effective number of species represents the number of species in a hypothetical community that has the same entropy (Shannon index, for q = 1) as the community at hand but completely even abundance. Similarly to the Shannon index, it weights species by their relative abundance, but unlike the Shannon index it is a true diversity metric (see Jost 2006 for details). Overall, wrack samples had higher diversity than larval samples although there was variation between sites and samples (Welch’s t-test, t = −5.02, *P* = 0.00008, Figure 2).

**Figure 2.**
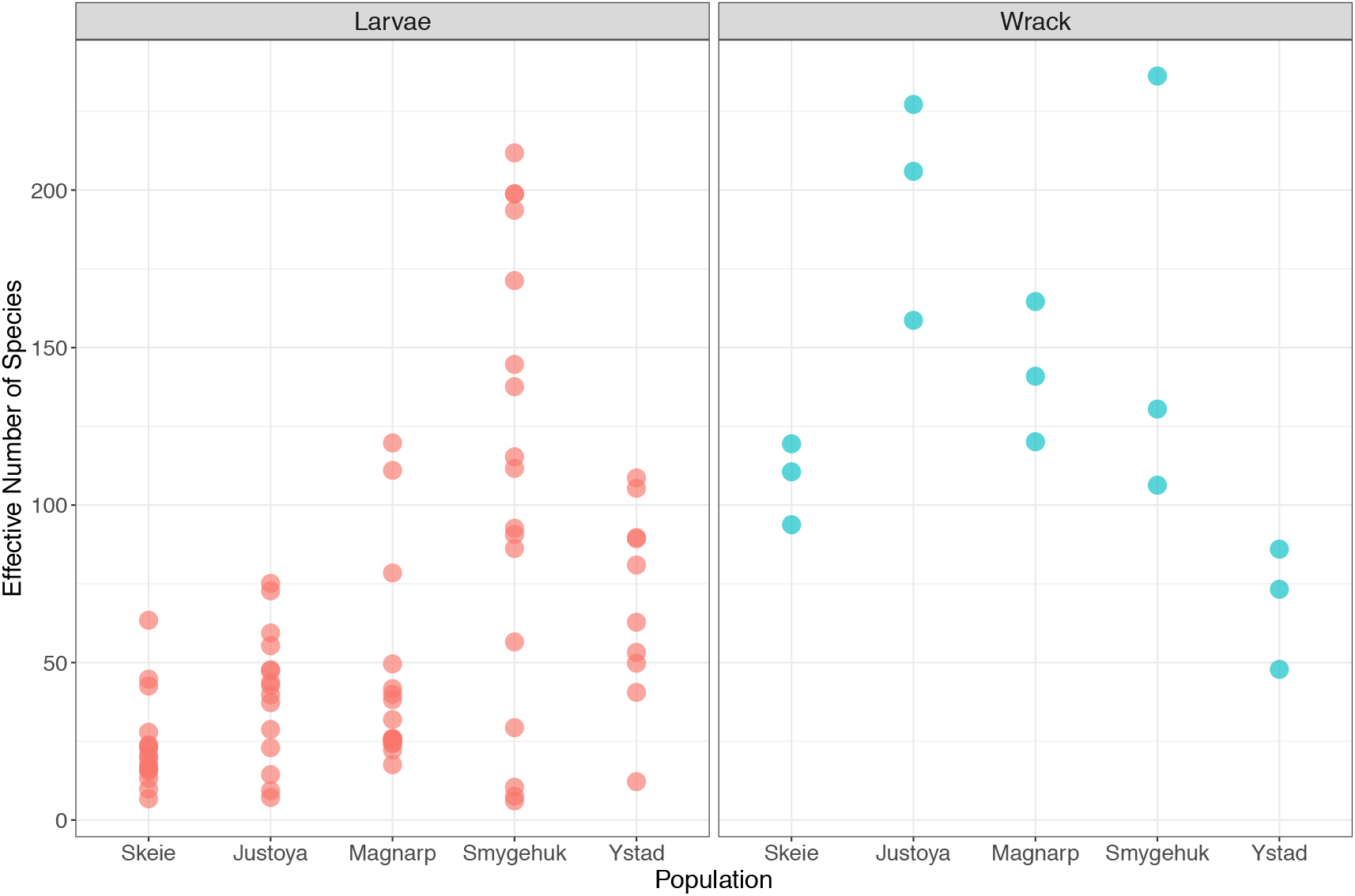
Effective number of species (e^shannon index^). Samples are colored by type, red-larvae, blue-wrack and labeled by site.

Phylum composition was similar among wrack sites but no significant core microbiome (i.e., set of common OTUs across samples) was found. We were able to assign 4,625 of our OTUs (99.4%) to 30 phyla. The most abundant phylum was Proteobacteria followed by Bacteroidota. There were clear and consistent differences in phylum composition between wrack and larvae (Figure 3, Table S3). Welch’s t-tests indicated that Verrucomicrobiota and Bacteroidota were more abundant in wrack samples while Actinobacteria and Proteobacteria were more abundant in larval samples (Table S3). However, there was no strong core wrack or larval microbiome: only 24 OTUs were found in all wrack samples and none comprised > 0.5% of the counts in even half of the samples. No OTU was found in all larval samples. There were also differences in phylum composition between Baltic and non-Baltic sites (Table S3). Spirochaeta, Desulfobacterota, Firmicutes, and Bacteroidota were more prevalent in the Baltic while Proteobacteria and Actinobacteria were more prevalent in the non-Baltic samples. As Bacteroidota and Proteobacteria were the only phyla significant in both larval vs. wrack and Baltic vs. non-Baltic analyses, we examined lower classification levels as well. There was no effect of sample type or location on Proteobacterial classes (data not shown). An examination of the order distribution within Bacteroidota revealed that the Flavobacteriales order was more prevalent in non-Baltic samples (t = −9.28, *P* = 3.042 e-13) while the Bacteroidales order was more dominant in the Baltic samples (Figure S3; t= 9.51, *P* = 2.161 e-12). There was no effect of sample type on class distribution within Bacteroidota. Full data prevalence and counts for all OTUs can be found in Table S4.

**Figure 3.**
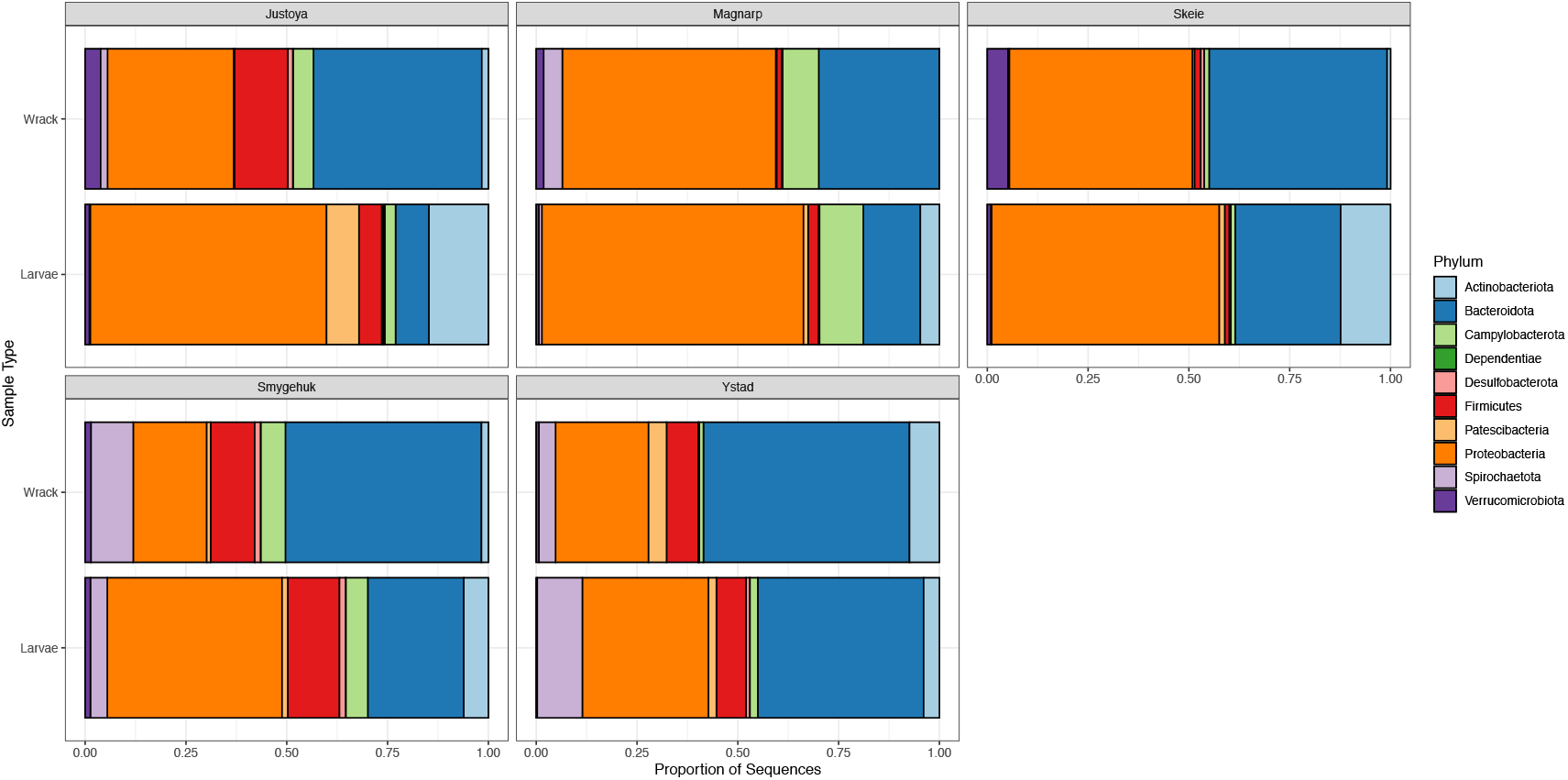
Taxonomic composition by site and sample type. All unassigned OTUs (30/4655) have been removed. All phyla that made up less than 1% of total counts have been removed.

Ordination of samples after the PhILR transformation revealed strong site level structure, but little effect of sample type. All sites except Ystad showed strong overlap between wrack and larval samples (Figure 4A,C). The strongest effect of site was an observed Baltic vs. non-Baltic split. The two Baltic sites (Ystad and Smygehuk) were separated from the rest of the sites along the first axis representing 27.7% of the variation and little differentiation among non-Baltic sites (Skeie, Justoya, and Magnarp) was observed (Figure 4C). This separation was statistically significant (F=25.27, df = 87, R^2^=0.225, *P* = 0.001, Adonis). When samples were plotted separately by type, some separation between the three non-Baltic sites was observed for wrack samples (Figure S5A). However, the same pattern did not hold true for larvae (Figure S5B).

**Figure 4.**
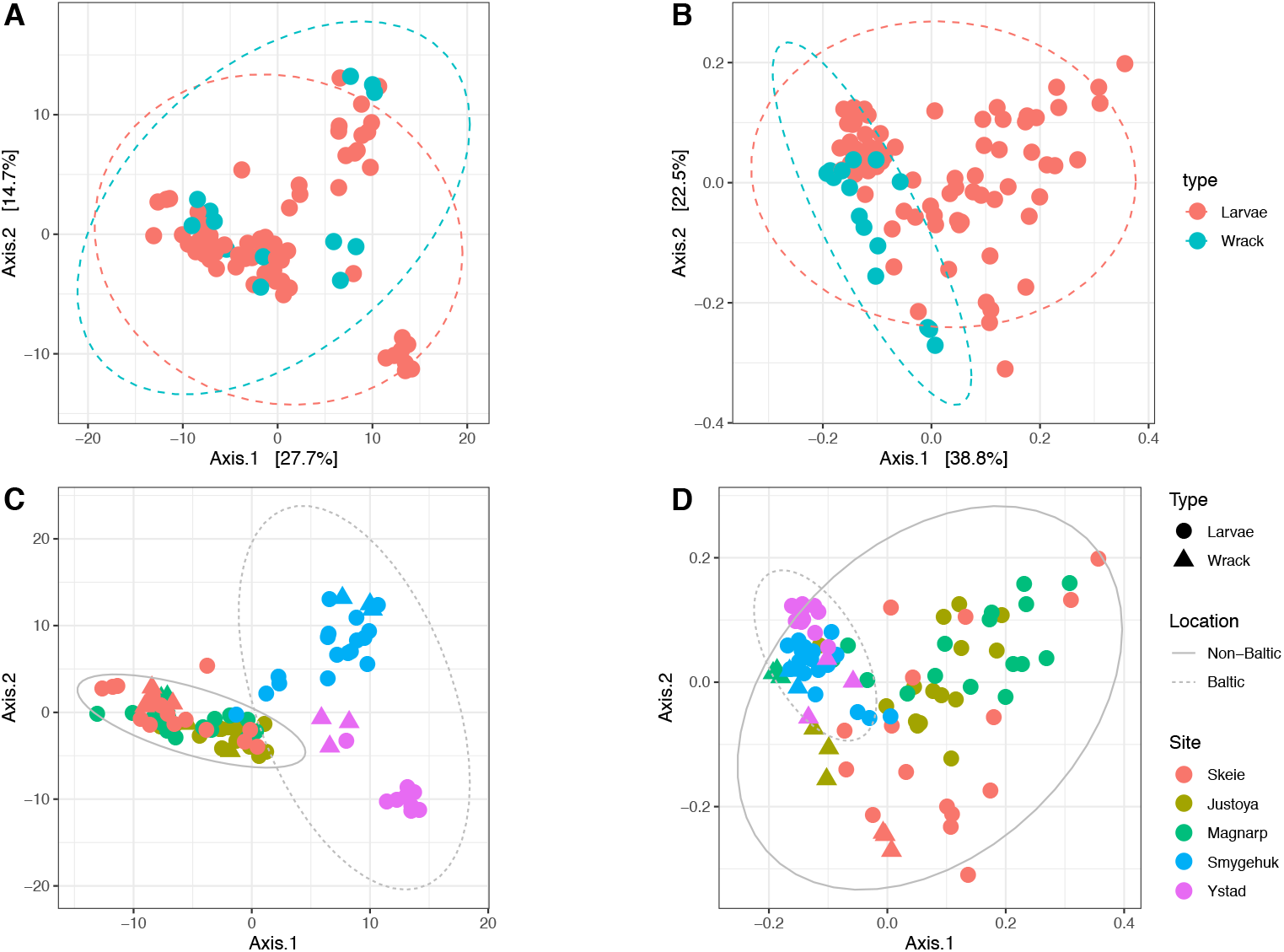
Baltic samples are compositionally and functionally separate. Ordination of samples after PhILR transformation (A,C) and ordination based on functional categories (B,D). Samples are colored by type (A,B) or by site (C,D). Sample type is indicated by shape in C and D. Dashed and solid lines indicate 95% ellipses.

The functional community profile indicated differences between sample types as well as a distinction between Baltic and non-Baltic sites. FAPROTAX identified chemoheterotrophy and fermentation as the most abundant categories (Figure S6). Unlike the phylum composition, there were no visible consistent functional differences between wrack and larval microbiomes. However, ordination based on functional categories revealed that larvae and wrack overlapped in functional space and that non-Baltic larvae were more variable (Figure 4B). A PERMDISP analysis confirmed this difference as significant (Table 2A). There were also severe changes in variance depending on the site (Figure 4D) with the two Baltic sites (Smygehuk and Ystad) tightly clustering together. A PERMDISP analysis confirmed that both site and location (e.g., Baltic vs. non-Baltic) were significant predictors of dispersion (Table 2B,C).

**Table 1.** Abiotic and biotic information for the collection sites. Note that wrack composition data for Justøya is not available.

| Population | Location | Mean Salinity* | Proportion Green Algae | Proportion Kelp | Proportion Fucoids | Proportion Red Algae |
| --- | --- | --- | --- | --- | --- | --- |
| Skeie | North Sea | 31.07 | 0.8 | 0.05 | 0 | 0.12 |
| Justøya | Skagerrak | 28.53 | N/A | N/A | N/A | N/A |
| Magnarp | Kattegat | 17.72 | 0 | 0 | 0.65 | 0.34 |
| Smygehuk | Baltic Sea | 7.95 | 0.05 | 0 | 0.45 | 0.5 |
| Ystad | Baltic Sea | 7.65 | 0.01 | 0 | 0.04 | 0.95 |

\*Based on yearly average from nearest monitoring station

**Table 2.** PERMDISP results for functional data from models testing A) type of sample, B) population, or C) location and D) Average distances to median for all groups calculated using the betadisper function in vegan

A. Type of Sample (Wrack or Larvae)
|  | Df | Sum of Squares | Mean Square | F | # of Permutations | Pr(>F) |
| --- | --- | --- | --- | --- | --- | --- |
| Type | 1 | 0.03301 | 0.033008 | 7.5968 | 5000 | 0.008998** |
| Residuals | 87 | 0.37801 | 0.004345 |  |  |  |

B. Population
|  | Df | Sum of Squares | Mean Square | F | # of Permutations | Pr(>F) |
| --- | --- | --- | --- | --- | --- | --- |
| Population | 4 | 0.26917 | 0.067292 | 14.997 | 5000 | 2e-04*** |
| Residuals | 84 | 0.37691 | 0.004487 |  |  |  |

C. Location (Baltic or non-Baltic)
|  | Df | Sum of Squares | Mean Square | F | # of Permutations | Pr(>F) |
| --- | --- | --- | --- | --- | --- | --- |
| Location | 1 | 0.2825 | 0.282501 | 97.144 | 5000 | 2e-04*** |
| Residuals | 87 | 0.253 | 0.002908 |  |  |  |

D. Average distance to Median
| Group | Average Distance to Median |
| --- | --- |
| Larvae | 0.215 |
| Wrack | 0.164 |
| Skeie Samples (larvae and wrack) | 0.217 |
| Justøya Samples (larvae and wrack) | 0.174 |
| Magnarp Samples (larvae and wrack) | 0.223 |
| Smygehuk Samples (larvae and wrack) | 0.103 |
| Ystad Samples (larvae and wrack) | 0.088 |
| Baltic Samples (larvae and wrack) | 0.111 |
| Non-Baltic Samples (larvae and wrack) | 0.228 |

We further investigated the differences between the Baltic and non-Baltic sites by identifying balances that distinguish the two groups. Balances are log-ratios of the geometric mean abundances of the two separate groups of taxa that descend from a node. We identified 8 significant balances (Figure 5) at different levels of taxonomy. The deepest node was the one separating Proteobacteria from the other phyla (n16). Indeed, Figure 3 shows a lower proportion of Proteobacteria in the Baltic samples compared to the non-Baltic samples and this was significant using a Welch’s t-test (Table S3). The other 7 nodes were at lower taxonomic levels (Figure 5B).

**Figure 5.**
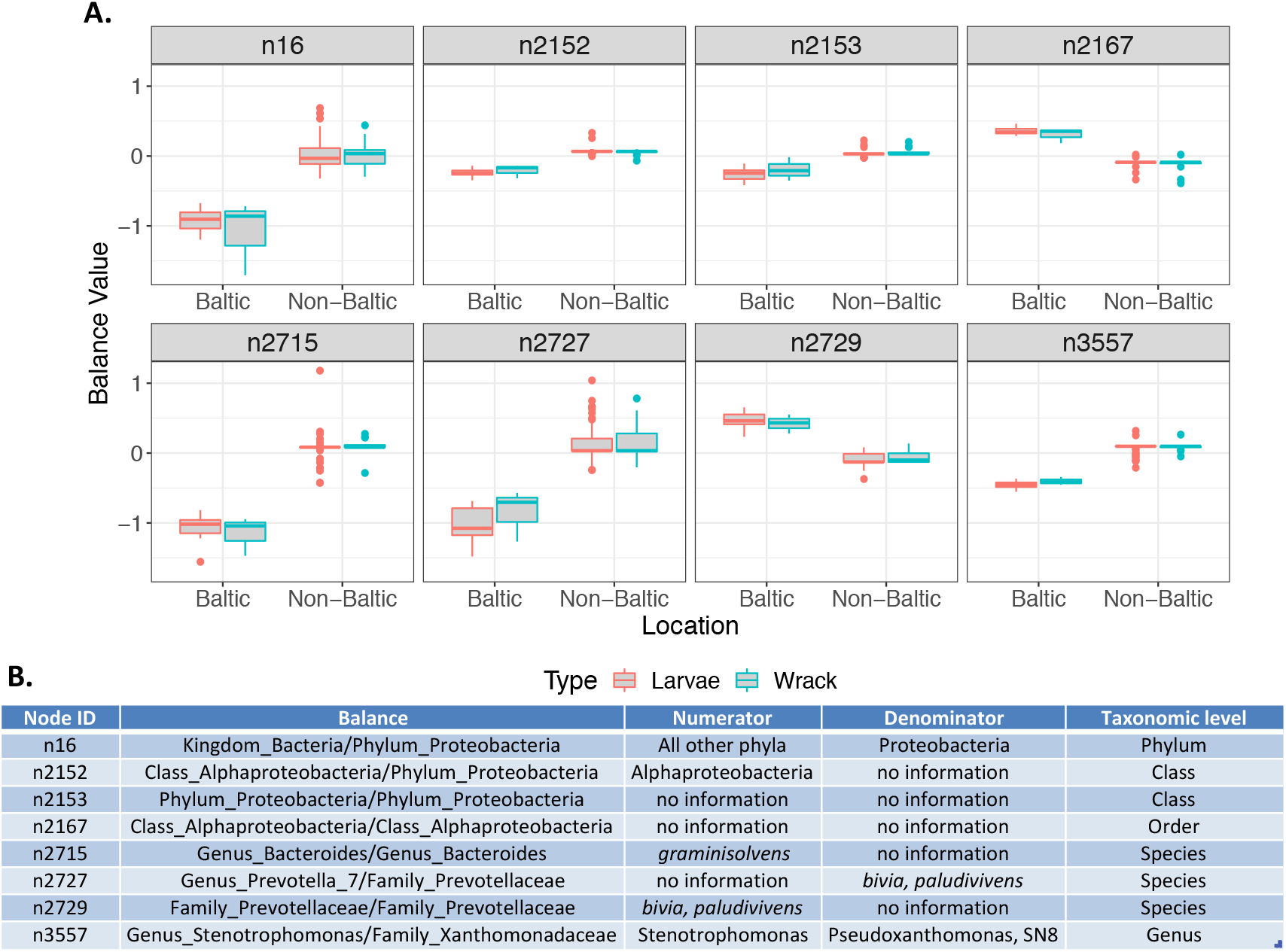
Balances separating Baltic and non-Baltic site. Balances represent the log-ratio of the geometric mean abundance of the two groups of taxa that descend from the given node. Internal nodes are numbered from base of the tree to the tips A. Boxplot showing the distribution of these balances for Baltic and non-Baltic wrack samples (blue) and larval samples (red). The number above corresponds to the node. B. Full data on each node with information on taxonomic groups that make up the numerator and denominator (when available) and the taxonomic level of the split.

There was no pattern between microbiome composition and genotype at *Cf-Inv(1)* in the *C. frigida* larvae. We were able to genotype 57/74 larval samples and found an over-abundance of heterozygotes (38/57) consistent with previous studies (Day, Dawe et al. 1983). Ordination after PhiLR transformation revealed no pattern based on genotype (Figure S7) although genotype sampling was not consistent between sites (Table S2).

## Discussion

Understanding the structure and function of complex microbial communities, and how they vary across space and time, is a major goal of microbial ecology. Here we report on the taxonomic and functional composition of the wrackbed microbiome as well as the microbiome of one common wrackbed bacterivore across the North Sea – Baltic Sea environmental transition zone. To our knowledge, this is the first report on the microbiome of deposited beach wrack at the OTU level. We observed strong overlap between sites following PhILR transformation (hereafter referred to as a coarse taxonomic scale) but were unable to identify a core microbiome. We also observed a separation between Baltic and non-Baltic sites in both bacterial taxonomic composition and functional variance, consistent with the environmental gradient in that zone although sampling of additional sites will be necessary to confirm this association. Below we discuss these patterns and their potential causes and consequences.

Sites differed at the OTU level but largely overlapped on a coarse scale. We were not able to find a strong core microbiome for the wrackbed; none of the 24 OTUs found in all wrack samples were highly prevalent (prevalence > 0.5%) in even half of the samples. This result may be partly explained by the idea of “functional redundancy” e.g., the idea that different metabolic functions can be performed by a wide range of taxa (Burke, Steinberg et al. 2011, Louca, Polz et al. 2018). This hypothesis is supported by the fact that the functional composition of the wrack samples overlapped substantially (Figure 4D), while the samples were completely separated in the Bray-Curtis NMDS (Figure S1) based on the species composition. However, the differences among non-Baltic sites were strongly reduced after the PhILR transformation (Figure 4C), indicating that the composition is similar on a coarse taxonomic scale even though the specific OTUs are different. In line with other studies, we propose that colonization of the wrackbed environment is likely a neutral process occurring via random dispersal (Hubbell 2006) with certain OTUs becoming abundant due to their functional properties (Burke, Steinberg et al. 2011, Louca, Polz et al. 2018). We were not able to determine the origin of these microbes (e.g., the path of colonization), although it is likely that a large number of them originated from the macroalgal microbiomes. However, a recent study on kelp detritus on the seafloor found that the microbiome shifted greatly as the kelp degraded (Brunet, de Bettignies et al. 2021), which may obscure the signal of origin.

The strong similarity between the non-Baltic sites on the coarse taxonomic scale is surprising as microbial communities are often highly dynamic (Louca, Jacques et al. 2016, Tully, Wheat et al. 2018). We suggest that the functional space of the wrackbed may be narrower than in other environments and that this may impose taxonomic constraints. Different functions are more redundant than others, for example, photoautotrophy is a more redundant function than sulfate respiration (Louca, Parfrey et al. 2016) and the wrackbed environment likely requires a large number of specific functions. For example, macroalgae contain secondary metabolites, such as phlorotannins which are polyphenolic compounds unique to brown seaweed (Glombitza and Kno 1992, Hierholtzer, Chatellard et al. 2013). These phlorotannins are often used as chemical defenses and are known inhibitors of anaerobic digestion systems (Chen, Cheng et al. 2008). Macroalgae also contain complex polysaccharides, the degradation of which requires highly specialized enzymes (Chauhan and Saxena 2016, Sichert, Corzett et al. 2020).

Our functional and compositional data indicate that these polysaccharide degraders are a dominant component of the wrackbed microbiome. Polysaccharides are major structural components in macroalgae and can comprise up to 50% of macroalgal biomass (Mabeau and Kloareg 1987). Our results show that chemoheterotrophy was the major functional category in all samples (Figure S6) and we found a compositional abundance of polysaccharide degraders. Members of the phylum Bacteroidota are the primary polysaccharide degraders in marine environments (Fernández-Gomez, Richter et al. 2013, Arnosti, Wietz et al. 2021), although Gammaproteobacteria (Sarmento, Morana et al. 2016), Planctomycetales (Reintjes, Arnosti et al. 2017), and Verrucomicrobia (Sichert, Corzett et al. 2020) are also known degraders. Bacteroidota was the second most abundant phylum with higher abundances in the wrackbed compared to the larvae and Proteobacteria was the most abundant phylum (Figure 3). Bacteroidota comprised 30-51% of the wrackbed microbiome compared to <10% of ocean water samples (Sunagawa, Coelho et al. 2015), although they are more common in macroalgal epiphytic bacterial communities (Florez, Camus et al. 2017) and are highly abundant on particulate organic matter (POM) (Fernández-Gomez, Richter et al. 2013).

As polysaccharide degraders are a major group in the wrackbed microbiome, we hypothesized that the specific polysaccharide composition of the macroalgal community nearby, and so also of the wrackbed, may be a major force shaping the microbial community. While all macroalgae contain polysaccharides, different groups of macroalgae contain different polysaccharides and the concentration can range from 4-76% of the dry weight (Kraan 2012). For example, in brown algae alginate can represent up to 60% of the total cell wall polysaccharides (Mabeau and Kloareg 1987). In red algae the most common polysaccharides are agarose and carrageenan (Popper, Michel et al. 2011) while porphyran is limited to the red alga *Porphyra* (Kraan 2012). Green algae contain sulphated polysaccharides such as Ulvan, which is a cell wall polysaccharide present in species of *Ulva* (Kidgell et al., 2019). Different carbohydrate-active enzymes (CAZymes) are needed to catabolize these compounds (Lombard, Golaconda Ramulu et al. 2014). A unique feature of *Bacteroidetes* genomes is that CAZymes are organized into polysaccharide utilization loci (PULs) that encode co-regulated enzyme and protein complexes for degradation of specific polysaccharides (Grondin, Tamura et al. 2017). Different species of *Bacteroidetes* contain different PULs specific to categories of polysaccharides (Grondin, Tamura et al. 2017). Closely related species of *Bacteroides* have been shown to be highly specialized on specific polysaccharide bonds, even to the point of neglecting the simple sugars these polysaccharides are built from (Martens *et al*., 2011). It has been seen that receptors on the outer membrane of *Bacteroides* specifically recognize complex carbohydrates and activate the appropriate PULs thus enabling the degradation of the triggering carbohydrates (Martens et al., 2011). There is a strong shift from brown and green to red algae between the non-Baltic and the Baltic sites that likely corresponds to a shift in wrackbed polysaccharide composition from ulvan, fucoidans, and alginate to agarose and carrageenan. This is accompanied by shifts in taxonomic composition of Bacteroidota. Three of the eight significant balances identified by our PhiLR analysis are within Bacteroidota (n2715, n2727, and n2729; Figure 5). Furthermore, an examination of the class distribution within Bacteroidota showed that the Flavobacteriales order was more prevalent in non-Baltic sites while the Bacteroidales order was more dominant in the Baltic sites (Figure S8). A metagenomic analysis looking at the frequency of different PULs in different wrackbeds will be necessary to formally test this link.

Compared to wrack samples, larval samples showed higher variation within site and lower site-specific signatures. Both the Bray-Curtis ordination (Figure S3) and the PhILR approach (Figure S5) showed that wrack samples tended to separate by site, but most larval samples did not. Only larvae from the Baltic sites (Ystad and Smygehuk) seemed to group by site. The functional analysis and PERMDISP also showed that larvae were more functionally variable than wrack samples, although seemingly with the exception of larvae from Ystad and Smygehuk (Figure 4B, Table 2A). Furthermore, no single ASV was present in every larval sample and although genotype sampling was highly uneven, none of the observed variance could be explained by genotype at the *Cf-Inv(1)* inversion (Figure S7). Some of this variation may be explained by experimental design: Wrackbeds are highly heterogeneous and contain many microhabitats, and the sampled larvae may have come from any number of these microhabitats whereas wrack samples were homogenized before sequencing, destroying microhabitat structure. Another potential source of variation is larval age. Larvae were taken directly from the field sites and were a variety of ages. Work in honeybees (Vojvodic, Rehan et al. 2013), leafworms and bollworms (Mason, St. Clair et al. 2020), and silkworms (Chen, Du et al. 2018) shows that there are strong shifts in larval microbiomes across instars.

Despite high variation in the larval samples, we observed consistent wrack-larval differences in the prevalence of Bacteroidota and Proteobacteria (Figure 3, Table S3). Our observed data include a combination of the “nutritional bacteria” that the larvae have ingested along with their own microbiome. As larval microbiomes do not directly match wrack microbiomes, it is clear that there is some level of selection in regards to either (1) which bacteria the larvae are eating and/or (2) which bacteria are colonizing and are getting established in the gut. The reduction in prevalence of the Bacteroidota is especially of note here as these are primary polysaccharide degraders. As polysaccharides are degraded, CAZymes as well as sugar oligomers and monomers can be released (Allison 2005, Teeling, Fuchs et al. 2012, Arnosti, Wietz et al. 2021). These can then be used by a wide variety of organisms including cheater or scavenger bacteria that cannot digest polysaccharides themselves. For example, in terrestrial ecosystems, detritivores show strong preferences for microbe digested substrates (Frainer, Jabiol et al. 2016). The relative rates by which *C. frigida* larvae consume bacteria (and if they preferentially consume certain bacteria), simple sugars, polysaccharides, and other compounds are still unknown, but it is possible that they preferentially take up simple sugars and other easily used nutrients. More detailed studies of wrackbed microbial ecology and the economics of polysaccharide degradation are clearly needed.

The observed Baltic-non-Baltic split coincides with numerous other physical and biological changes occurring over the same spatial gradient (Snoeijs-Leijonmalm, Schubert et al. 2017). Perhaps the most powerful physical driver of biological systems that varies across this gradient is salinity, which ranges from 8-10 psu in the Baltic up to >30 psu in the North Sea and can vary seasonally (Møller Nielsen, Paulino et al. 2016). This and other biological gradients can have powerful effects on the marine life of the region (Pearson, Kautsky et al. 2000), resulting in observed genetic breaks in many species (Johannesson, Le Moan et al. 2020), a pattern which holds true in our data. However, we note that we only sampled five sites along the gradient and this pattern might change with more intensive sampling. Despite this caveat the observed pattern is consistent with dependence of the wrackbed environment on the seaweeds that grow nearby. Still it is remarkable that these differences are sustained through multiple trophic linkages and spatial subsidy events as the seaweed is washed ashore and degrades.

## Conclusion

Wrackbeds are biogeochemical hotspots where a combination of microbes and grazers degrade stranded seaweed and provide the base of a complex food web. Polysaccharides make up the major component of algal carbon and our results indicate that the wrackbed microbiome is specialized for polysaccharide degradation. Furthermore, the microbiome composition potentially alters based on the polysaccharides present. This change of microbiome composition co-occurs with a strong change over a natural marine environmental transition zone (the entrance of the Baltic Sea), which may be directly influenced by the changes in abiotic factors like salinity, or indirectly through the changing seaweed community, which is controlled by those abiotic factors. This shift carries up through tropic levels to the microbiome of seaweed fly larvae although larvae were more variable than the wrackbed itself. However, no connection between genotype at the *Cf-Inv(1)* inversion and larval microbiome was found, indicating that the wrackbed microbiome may not be a driver of selection in this species. The microbial food web of the wrackbed is potentially very complex, but studies of wrackbeds are currently in their infancy and the diverse roles of the various bacterial groups remain a black box at present.

## Acknowledgements

E.L.B. was supported by a Marie Skłodowska-Curie fellowship 704920 – ADAPTIVE INVERSIONS and gratefully acknowledges funding from Helge Ax:son Johnsons Stiftelse. Amplicon analysis was enabled by resources provided by the Swedish National Infrastructure for Computing (SNIC) partially funded by the Swedish Research Council through grant agreement no. 2018-05973. Additional funding was provided by the Swedish Research Councils VR and Formas through the Linnaeus Centre for Marine Evolutionary Biology, CeMEB.

## Supplemental Figure Legends

<u>Figure S1</u> - Bray-Curtis ordination of the wrack samples. Samples are colored by site. Samples are named site-biological sample-technical replicate. Note the spurious Skeie sample (Skeie-3-2).

<u>Figure S2</u> - Histogram and density plot of prevalence of ASVs among all samples.

<u>Figure S3</u> - Bray-Curtis ordination of all samples. Samples are colored by site.

<u>Figure S4</u> - Rarefaction curves for (A) Wrack samples and (B) Larval samples. Curves were made using the rarecurve function in the vegan package (Dixon 2003) with a step size of 50.

<u>Figure S5</u> - Separate ordination of wrack (A) and larval (B) samples after PhiLR ordination. Samples are colored by site.

<u>Figure S6</u> - Functional composition of samples by site and sample type. Only annotated OTUs are shown here (24%). Groups that had a Jaccard similarity index of >0.75 with another group have been removed.

<u>Figure S7</u> - Ordination of larval samples after PhiLR ordination colored by genotype.

<u>Figure S8</u> - Class composition of Bacteroidota by site and sample type. All classes that made up less than 1% of total counts have been removed.

## Supplemental Tables

<u>Table S1</u> - Detailed seaweed composition information for the different sites. Note that composition information for Justoya is unavailable.

<u>Table S2</u> - Full information on all samples.

<u>Table S3</u> - Welch’s t-tests for differences in phylum abundance. Bold indicates significant p-values. Note that calculation of df includes standard deviations.

<u>Table S4</u> - Full ASV table with taxonomic information and prevalence statistics.

