## Supplementary figures and images for "A metagenomic analysis of the wrackbed microbiome indicates a phylogeographic break along the North Sea - Baltic Sea transition zone"

### Figure S1

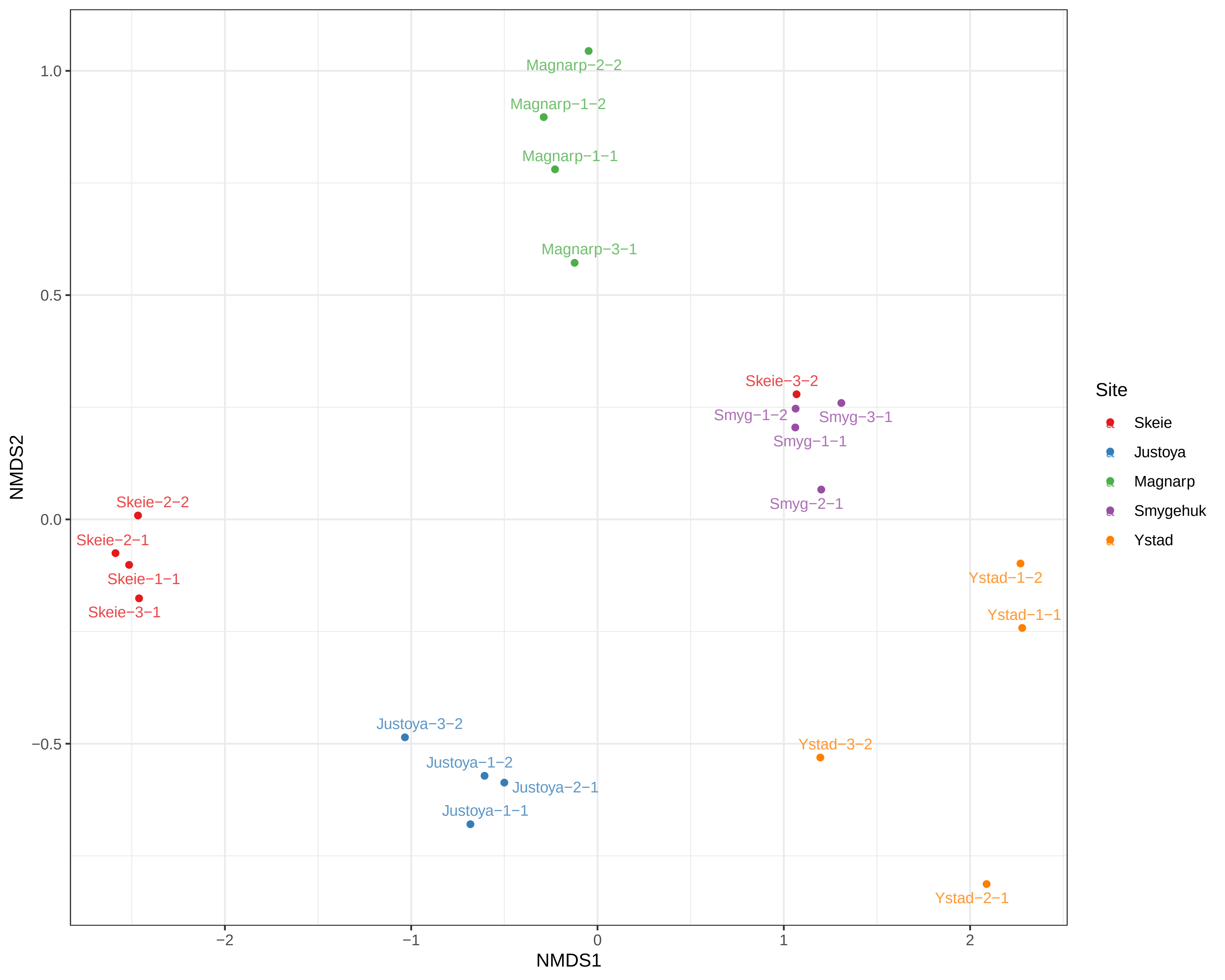

### Figure S2

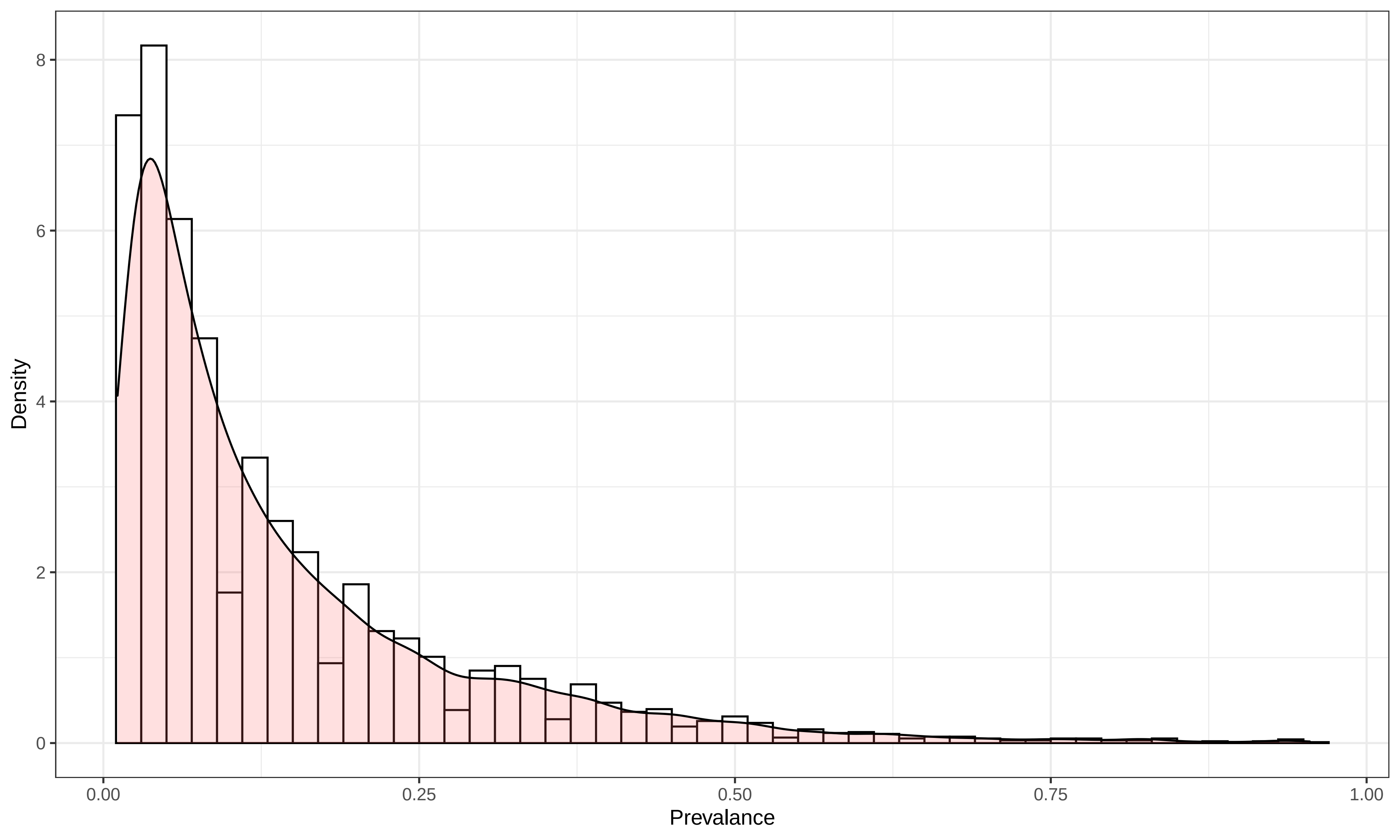

### Figure S3

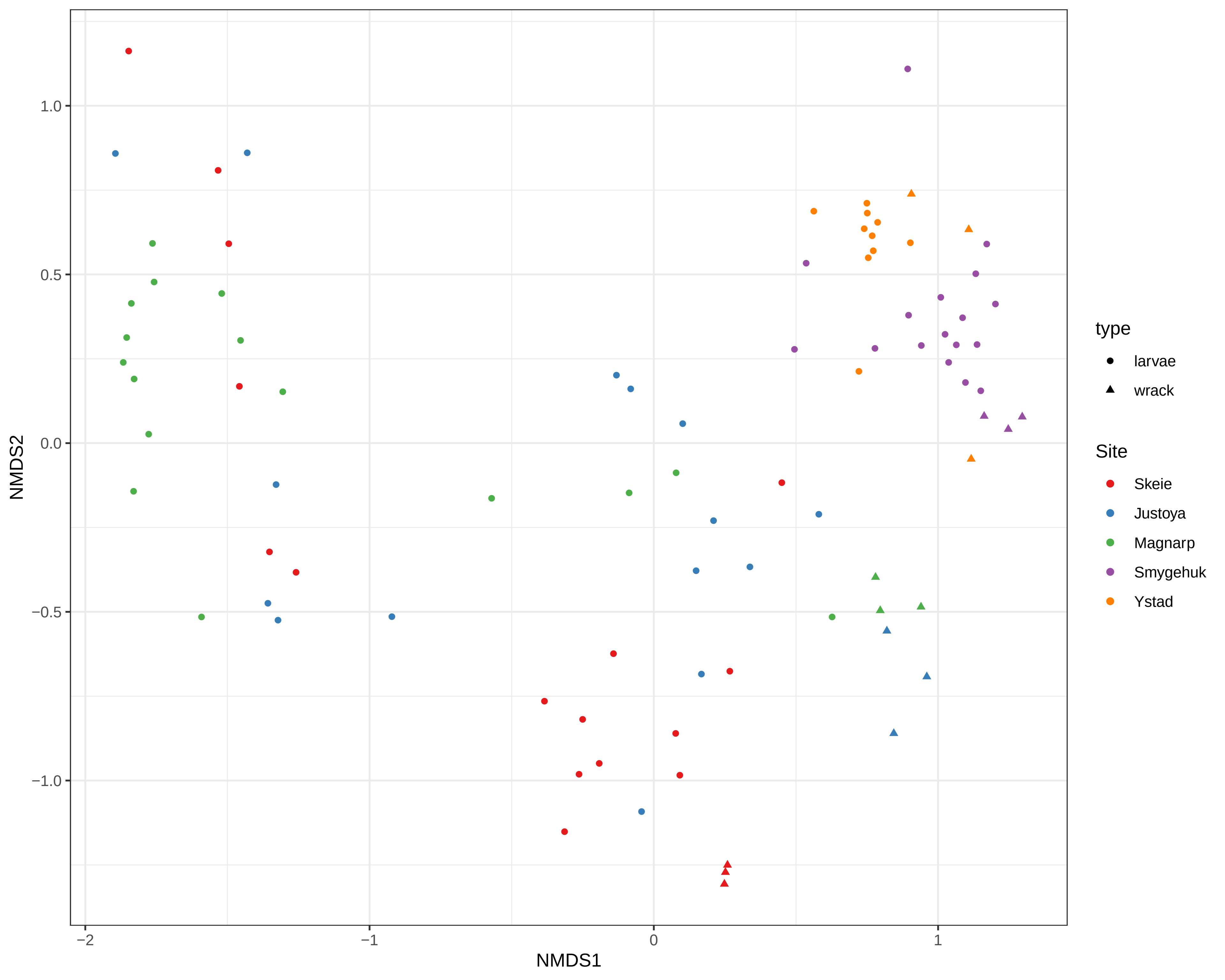

### Figure S4

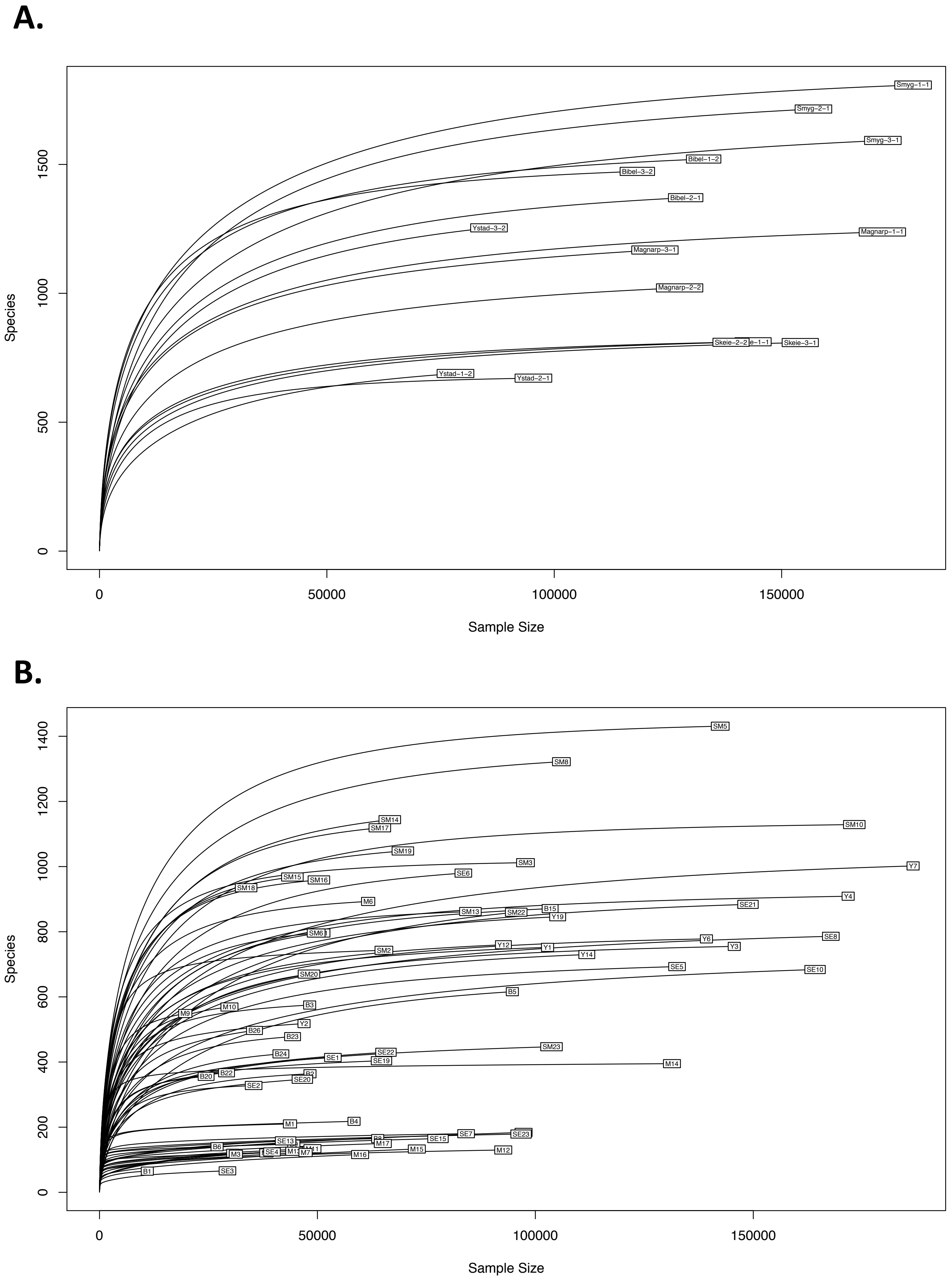

### Figure S5

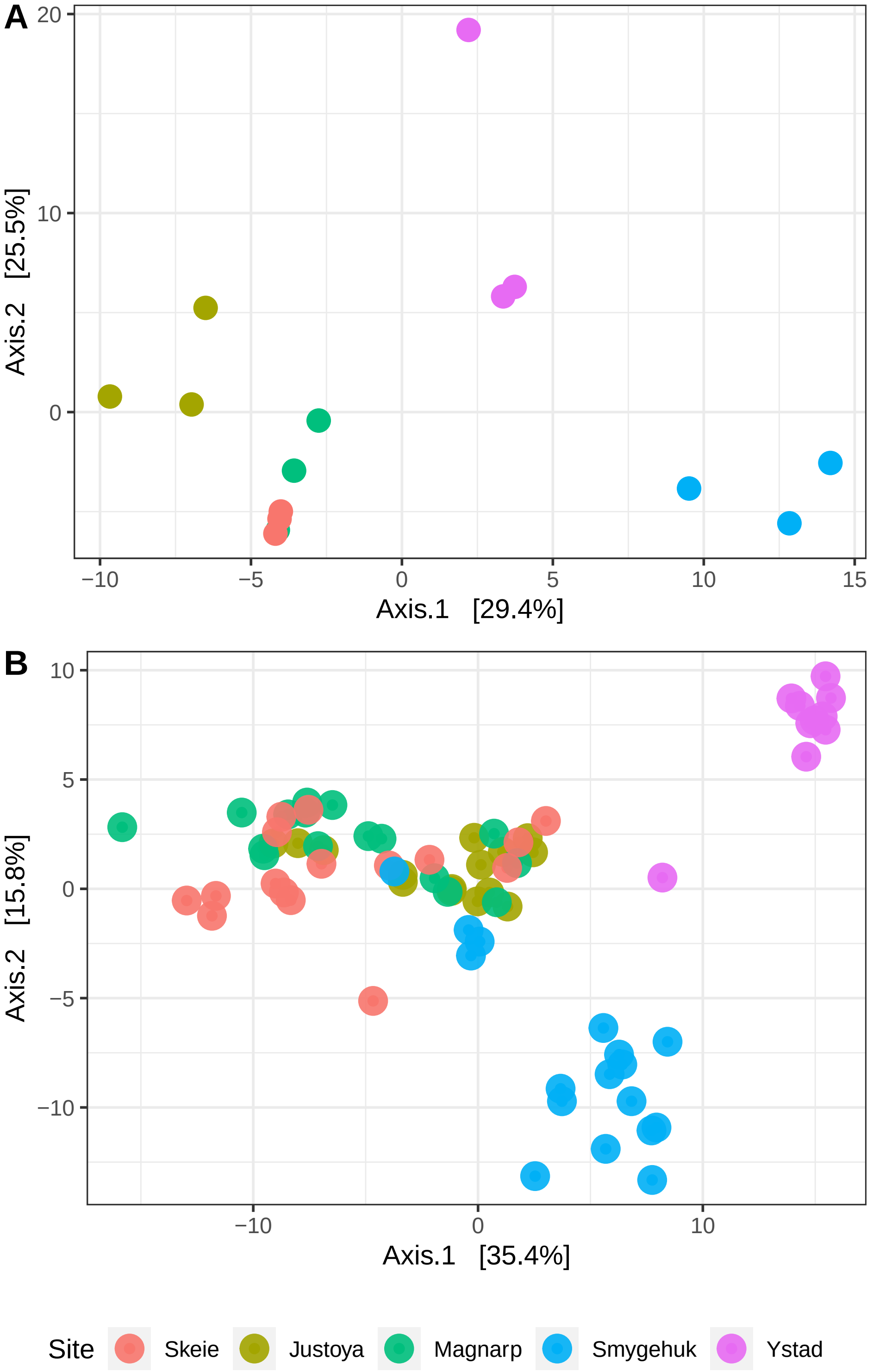

### Figure S6

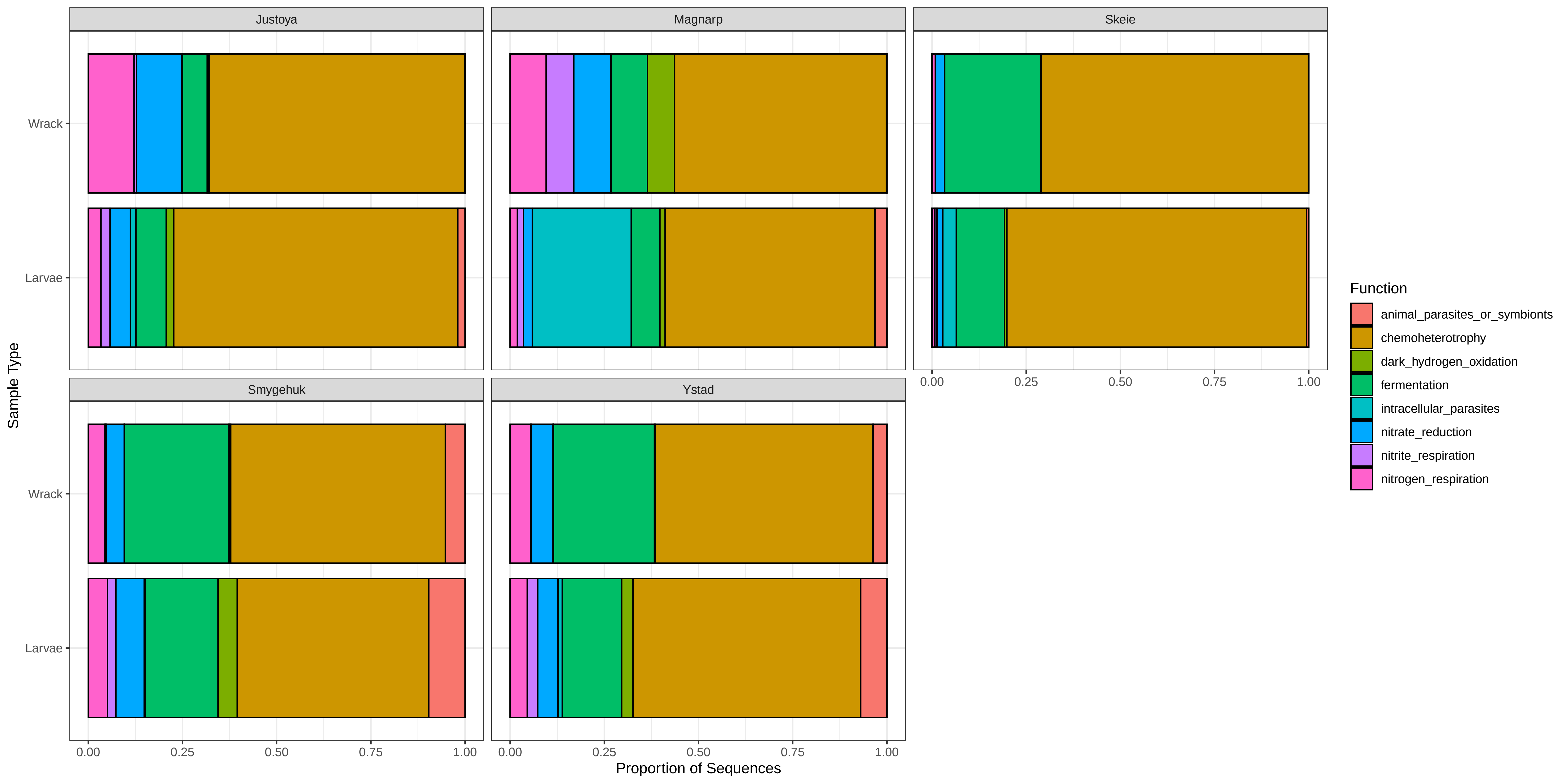

### Figure S7

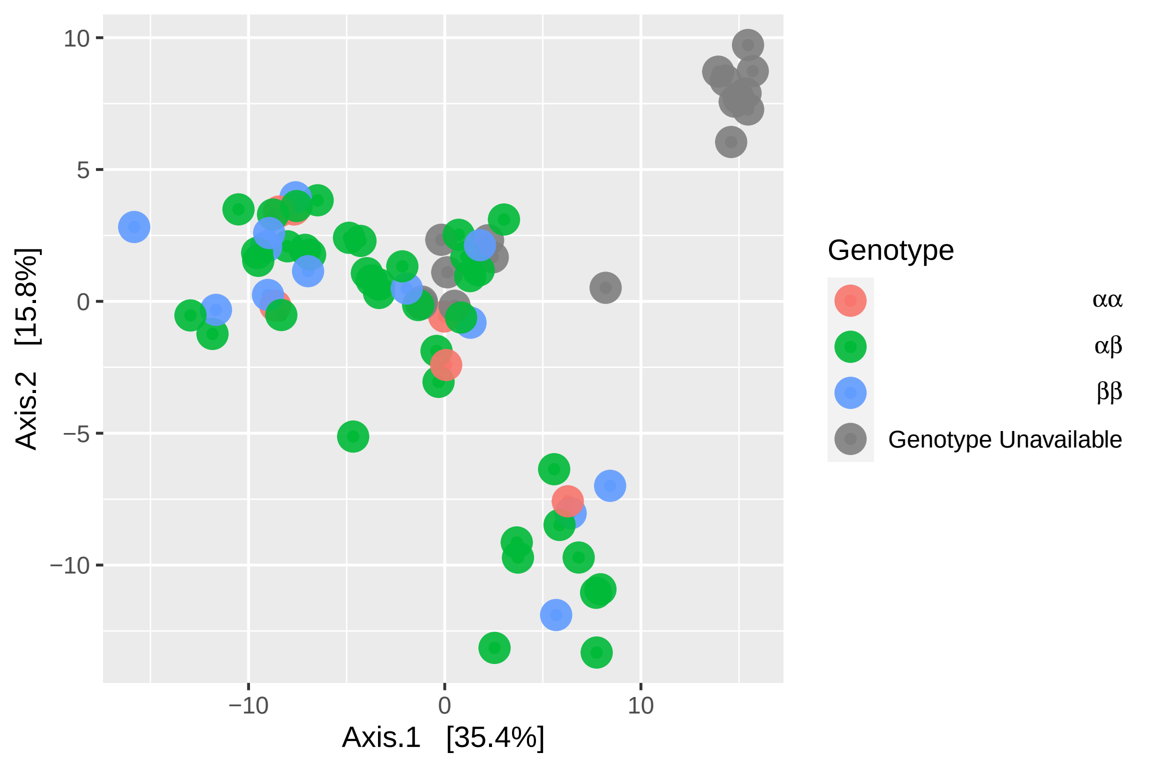

### Figure S8

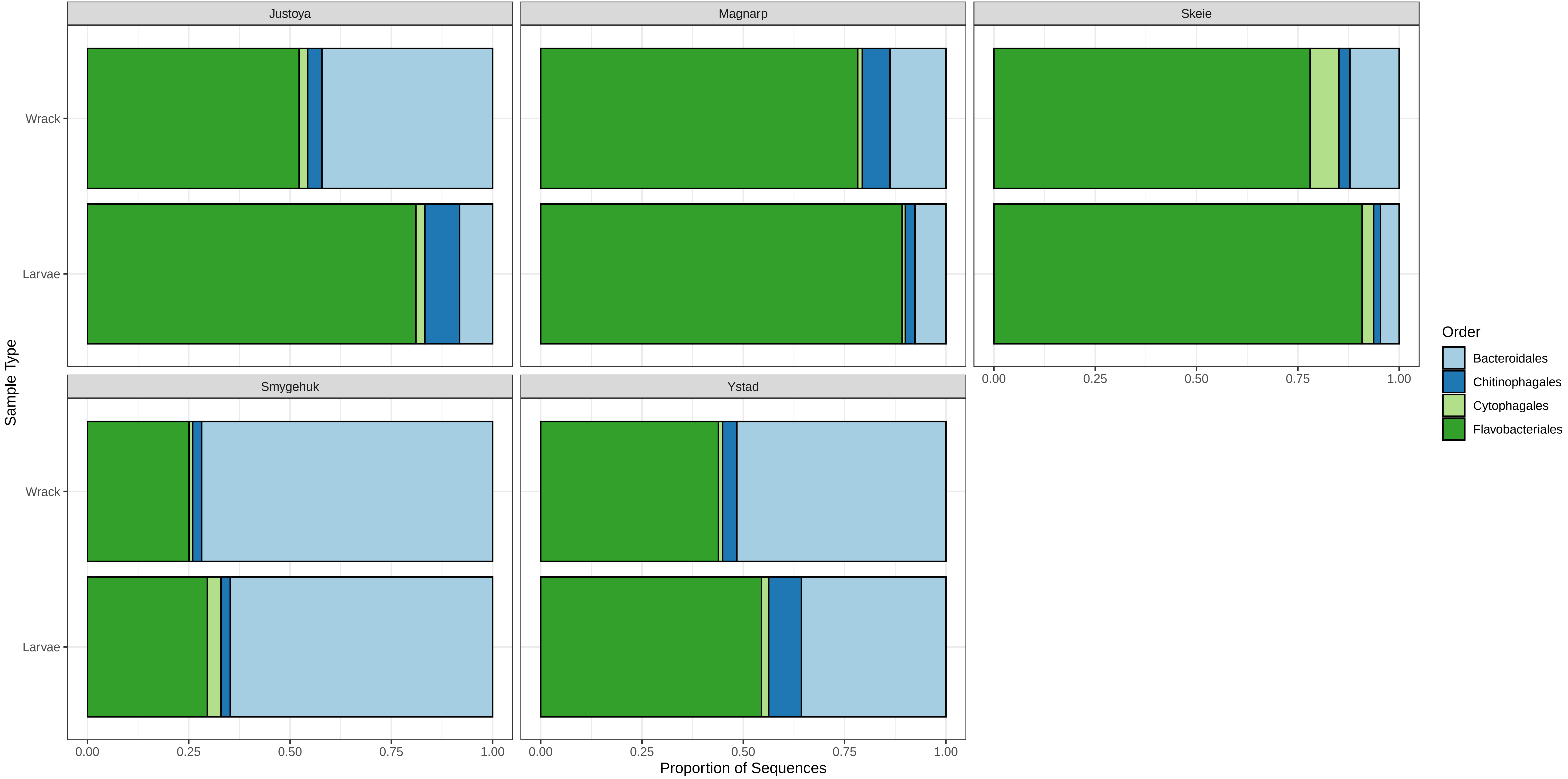
